# Time is Confidence: Monetary Incentives Metacognitive Profile on Duration Judgment

**DOI:** 10.1101/2023.11.10.566595

**Authors:** Mitra Taghizadeh Sarabi, Eckart Zimmermann

## Abstract

The question we addressed in the current study is whether the mere prospect of monetary reward affects subjective time perception. To test this question, we collected trail-based confidence reports in a task in which subjects made categorical decisions about probe durations relative to the reference duration. When there was a potential to gain monetary reward, the duration was perceived to be longer than in the neutral condition, and confidence, which reflects the perceived probability of being correct, was higher in the reward condition than in the neutral condition. We found that confidence influences the sense of time in different individuals: subjects with high-confidence reported that they perceived the duration signaled by the monetary gain condition as longer than subjects with low-confidence. Our results showed that only high-confidence individuals overestimated the monetary gain context. Finally, we found a negative relationship between confidence and time perception, and that confidence bias at the maximum uncertainty duration of 450 ms is predictive of time perception. Taken together, the current study demonstrates that subjective measure of the confidence profile caused overestimation of time rather than by the outcome valence of reward expectancy.

## Introduction

Monetary reward is a well-known motivator for individuals to perform tasks or achieve goals^1^. It has also been recognized as an extrinsic incentive that can influence decision making and time perception^2^. Time perception refers to the subjective experience of time^3^, including its duration^4^, speed^5^, and sense of time^6–8^. Confidence, on the other hand, relates to an individual-level of self-assurance^9^ in their abilities and decision making^10–12^. It is a multifaceted construct that is influenced by a range of factors, including self-esteem, experience, and motivation^13^.

A number of studies have shown that monetary rewards can affect the perception of time^2,14,15^. For example, cognitive-behavioral findings in healthy human subjects showed that the expectation of monetary reward could alter the subjective perception of duration in short time intervals. One piece of evidence showed that when a high amount of money was associated with an oddball disc, the perception of the oddball’s duration was overestimated compared to an oddball associated with a low amount of money or no money^15^. Notably, when a monetary reward was presented before the oddball and not by the oddball itself, the perception of duration remained unaffected. However, it has also been reported that when there is a potential to win money, the duration is perceived as longer than in loss or neutral conditions. In other words, cuing a monetary reward prior to a duration judgment task distorts time, causing it to be overestimated compared to the reference duration^16^. Given the aforementioned different monetary reward conditions, a considerable body of literature generally associates attentional^17–22^ or intentional^23–26^ resources with monetary gain, which causes a longer perception of duration.

Another stream of literature has shown that monetary rewards can affect human confidence^27^. The expectation of a monetary reward can increase individuals confidence in their ability to perform a task, leading to better performance. For example, a study showed that confidence was behind the prospect of monetary gain in reinforcement-learning strategies^27^. In this study, participants’ learning strategies differed between seeking gain and avoiding loss, with the former showing a higher confidence score. Neurophysiology studies also reported that monkeys did not select the sure target based on the difficulty of the stimulus but rather based on a sensation of uncertainty on each trial, indicating that the source of information about difficulty is not solely controlled by stimulus characteristics but also by internal variability that affects how reliable the evidence is to the decision-maker^28^. Also, neural recordings in rats proposed that a confidence estimate might be a basic and pervasive element of decision making, and the likelihood of a successful trial outcome may theoretically be calculated using a subjective indicator of decision-making confidence^29^.

Given the importance of confidence in decision making, however, the relationship between confidence and time perception is lacking in the literature^30,31^. The current study investigated whether differences in confidence determine how subjects perceive time in a monetary context. To estimate whether confidence distorts duration judgments, physically and mentally healthy individuals were tested. Verifying previous findings^15,16^, we demonstrated that the perceived duration of the monetary gain condition was perceived as longer than the neutral condition and that confidence, as the perceived likelihood of being correct, was higher in a monetary gain scenario than in neutral and loss scenarios. We found that individual differences in confidence influenced duration judgments. For almost half of the subjects, perceived duration was influenced by confidence but not for the other half. Subjects with high-confidence perceived the monetary gain condition as longer than the neutral condition, whereas subjects with low-confidence did not. We also found a correlation between time perception and confidence level, where high-confidence individuals were more engaged by the monetary gain contexts, which may lead to duration overestimation. Moreover, linear regression analysis revealed a stronger relationship between confidence of 450 ms and time perception and that confidence of maximum uncertainty at the duration of 450 ms is predictive of the perceived time in the monetary gain condition, and the more confident subjects are in their longer responses at the 450 ms the time will be more overestimated.

## METHODS

### Participants

Twenty-four healthy volunteers (18 females, aged 19–37 years, mean = 23.01, all right-handed, all with normal or corrected-to-normal vision) were recruited from the campus of the Heinrich-Heine-University Düsseldorf. None had a history of psychiatric or neurological disease, and none were taking any drugs or medication at the time of testing. The sample size was based on previous similar studies^14^. Written informed consent was obtained from all subjects prior to participation in the study. Subjects were tested in exchange for monetary compensation (€10 per hour) or course credits, and were additionally compensated based on their performance in a randomly selected session. This study was approved by the local ethics committee of the Department of Psychology, Heinrich-Heine-University, Düsseldorf in accordance with the Declaration of Helsinki.

### Stimuli

Adapted from the monetary incentive delay (MID) task^32^, the monetary incentive was an ecological picture of 50 cents (€0.5) outlined in blue, red, or gray colors signaled gain, loss, and no monetary outcome, respectively (see Fig. 1). For half of the participants, the gain cue was presented in blue and the loss in red; for the other half, the colors were reversed. The neutral cue was presented in gray for all participants. A white circle sized (1.93º) was used as both references and probe stimulus. All stimuli were presented on screen (LCD, 27 inches, 240 Hz NVIDIA’s G-Sync, Acer XB272) with a resolution of 1920 × 1080 pixel and a refresh rate of 60 Hz. Stimuli were presented with Presentation V2.4. Each stimulus was centered on the screen with a homogeneous dark gray background (see Fig. 1).

**Fig. 1.**
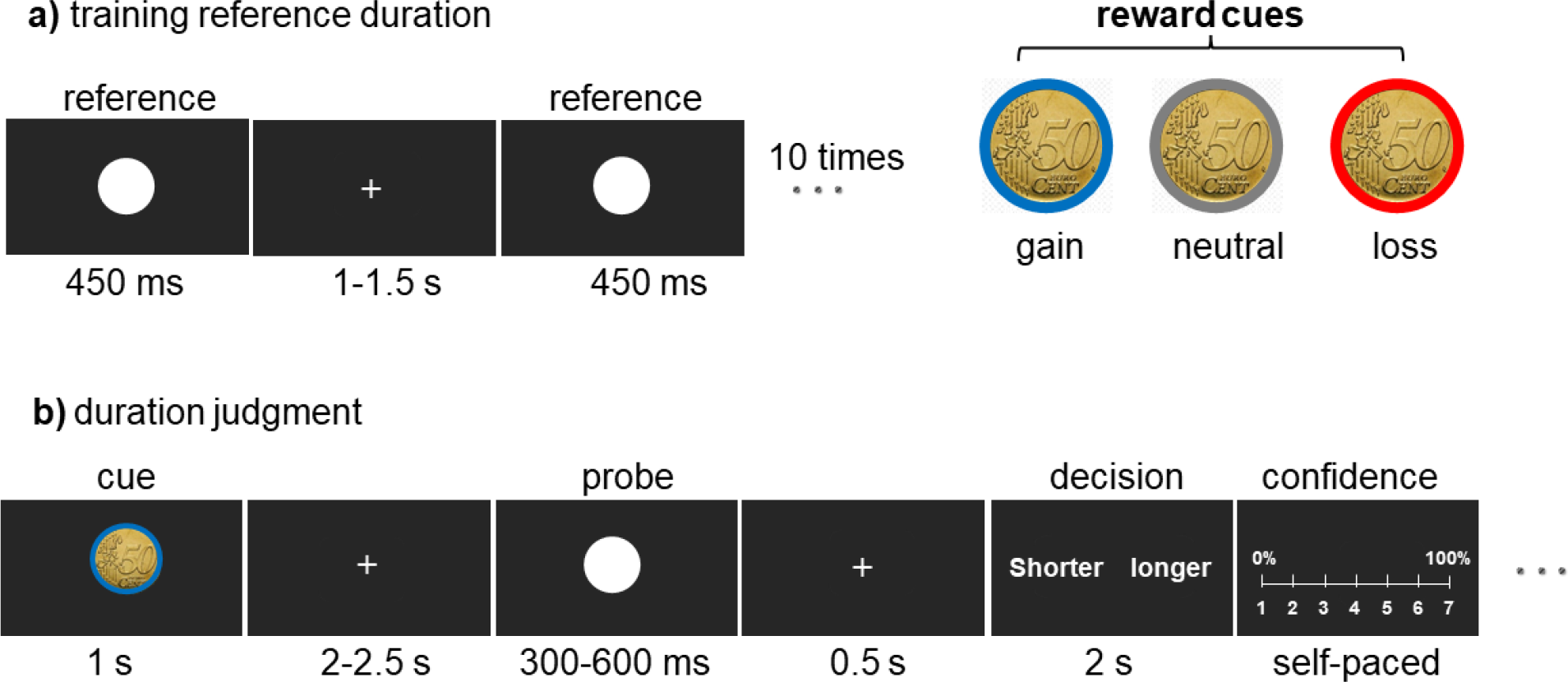
Experiment set-up. **a)** Training reference duration (approximately 2 min). After a 5 s fixation cross, participants viewed a flashing white circle ten times with a random Inter-trial interval (1-1.5 s) to learn the reference duration of (450 ms). **b)** An example trial of the duration judgment task. The probe circle was selected from 300, 350, 400, 450, 500, 550, or 600 ms, followed by a monetary incentive cue of 1 s in either the blue, red, or gray color with a random fixation (2-2.5 s) in between. Subjects responded to the decision probe within the 2 s time limit. They pressed a mouse button to indicate that the perceived probe duration was longer or shorter than the reference duration. Each trial was scored on a Likert scale from 1 (0% confident) to 7 (100% confident) to obtain a confidence score.

### Experimental procedure and design

#### Instruction

Participants were given instructions for the entire experiment while sitting 70 cm away from the computer screen in a dimly lit room. To maintain participants’ motivation throughout the experiment, they were told that the reward outcome would be determined cumulatively according to their performance and that the final reward would be based on only one randomly selected session. All subjects completed a practice version of the task for approximately 2 min or until they demonstrated proficiency in the task before beginning the first session.

#### Task Design

Participants performed a prospective duration judgment^33^ experiment divided into five sessions, all performed on a single day and separated by short rest periods. In each session, subjects first learned a reference duration (450 ms) presented by a flash of a white circle (see Fig. 1.a) ten times with a variable interval (1-1.5 s). The training session was performed before each main session. Immediately after the training reference period, the main session began, consisting of 63 trials (∼10 min). During each trial, participants saw one of the three €0.5 monetary incentive cues (red, blue, or gray) for (1 s). After a random fixation delay of (2-2.5 s), the probe stimulus flashed for a variable duration drawn from 6 equiprobable durations (300-600 ms in 50 ms steps). After a (0.5 s) fixation delay, a decision screen appeared. Participants had (2 s) to respond whether they perceived the probe duration shorter or longer than the reference (two-alternative forced-choice task (Fig. 1.b). They registered their responses by clicking the left or right mouse button with their index and middle fingers, and the experiment continued as soon as participants pressed one of the two response buttons within the response time. A self-paced 1-7 Likert scale was added at the end of each trial, and participants were asked to indicate their decision confidence by moving the mouse from 1 (0%, not at all certain) to 7 (100%, definitely certain). Finally, the probe screen was replaced by a central fixation cross that jittered randomly in the range (3-7 s). Subjects were also instructed to fixate the fixation point throughout the sessions. Note that only the first trial in each session started after a fixed (5 s) fixation delay. No feedback was provided for participants to avoid stress^34^ and learning effects on memory^35,36^. A single value was used for reward and punishment conditions to avoid the parametric effect of the motivational effect associated with the size of a potential reward^37,38^.

#### Supplementary information

The placement of probe texts (longer or shorter) was counterbalanced across trials. Onset times, response accuracy, and reaction times were recorded using Presentation V2.4. Reward outcomes were determined by task performance on each trial; subjects gained 0.5 cents for a correct response in the gain condition and lost 0.5 cents for an incorrect or missed response in the loss condition. The neutral condition had no effect on reward outcomes. In the practice version, only the easy target durations (300 and 600 ms) were tested ten times (five times each) in a random order. In the main sessions, all target durations were tested equally (9 times per session). Participants were informed that only the target duration would vary in the display, while the incentives would be displayed for a fixed duration throughout the experiment. Each participant completed 315 trials. The duration of the experiment was approximately 60 min.

### Analysis and psychometric function

We analyzed the proportion that participant reported the probe stimuli lasting longer than the standard. From this data, we estimated the psychometric curve for each reward condition separately implemented in *quickpsy*^39^ (http://dlinares.org/quickpsy.html). The point of subjective equality (*PSE*: the temporal duration at which subjects felt equal to the reference temporal duration, i.e., the 50% probability to report the prob lasting longer) and slope were calculated for each condition. Each participant’s *PSE* and slope for each reward condition (gain, loss, and neutral) were used for statistical significance testing using paired-samples t-tests once normality was demonstrated (Shapiro–Wilks test). If the assumption of normality was violated, a nonparametric Wilcoxon signed-rank test was used. Note that in these cases, the median (Mdn) is reported instead of the mean and standard deviation. All statistics were calculated on the basis of 95% confidence interval (95% CI).

## Results

### Effect of reward on time perception

To investigate whether reward gain influences time perception, we compared the *PSE*s between reward conditions. The *PSE* gain had lower values for outcome (*M* = .46, SD = .03) than the *PSE* neutral (*M* = .47, SD = .03). A paired-samples t test showed that this difference was statistically significant; *t*(17) = -2.76, *p* = .0067 (one-tailed), CI [-.018, - .002], Cohen’s d_av_ = .32 (see Fig. 2.a). A paired-samples t test revealed no statistically significant differences between (loss vs. neutral); (*M* = .47 vs. *M* = .47),*t*(17) = -1.33, *p* = .201, CI [-. 018, .004], Cohen’s d_av_ = .24, as well as between (gain vs. loss); (*M* = .46 vs. *M* = .47), *t*(17) = -.68, *p* = .505, CI [-.013, .007], Cohen’s d_av_ = .11, indicating that participants perceived only the monetary gain condition to last longer than the neutral condition. A Wilcoxon signed-rank test revealed no differences in the slope values between (gain vs. neutral), (*Mdn* = 106.195 vs. *Mdn* = 110.678), *z* = -1.33, *p* = .196, *r* = - .357, between (gain vs. loss), (*Mdn* = 106.195 vs. *Mdn* = 110.678). loss), (*Mdn* = 106.195 vs. *Mdn* = 103.397), z = -1.023, *p* = .325, *r* = -.275, and between (loss vs. neutral), (*Mdn* = 103.397 vs. *Mdn* = 110.678), *z* = -1.023, *p* = .325, *r* = -.275, (see Fig. 2.a)

**Figure 2.**
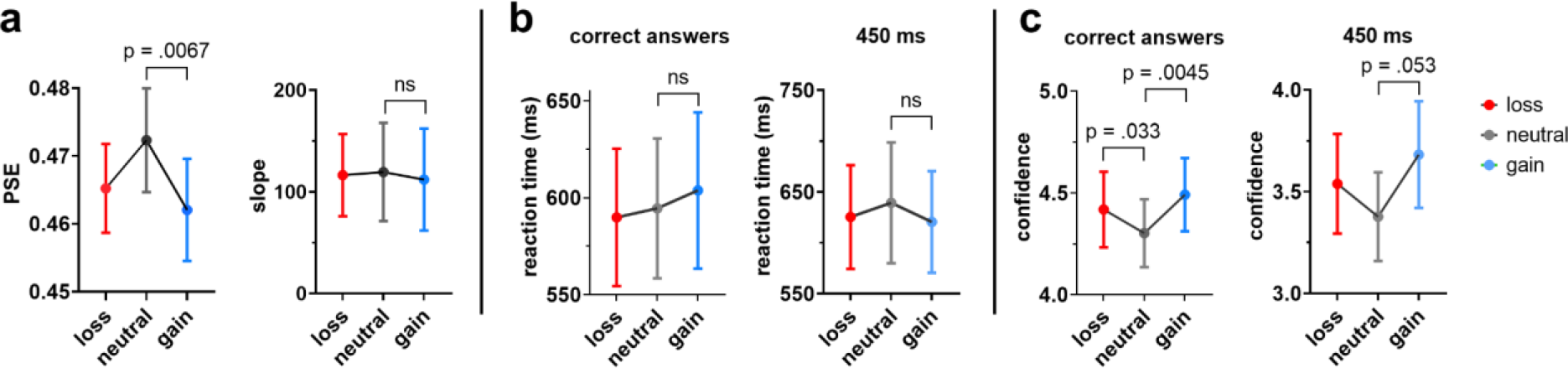
**a)** *PSE* for each reward condition shows a significant difference between gain and neutral (left), slope values of the fitted psychometric function (right). **b)** Mean values of reaction time across all probe durations except 450 ms (left), mean values of reaction time that perceived longer for 450 ms (right). **c)** Mean confidence scores for all probe durations except 450 ms (left), mean values of confidence ratings perceived longer only for 450 ms (right). (Note: all bar graphs show mean and standard error, and 450 ms is plotted separately since it is equal to the reference duration and no correct response is given for this duration).

#### Reaction time of perception (RTP)

We averaged reaction time scores of correct answers for each reward condition (gain, loss, and neutral) across six probe durations (300, 350, 400, 500, 550, 600 ms), (see Fig. 2. b, left) and reaction time of answers that subjects perceived to be longer on probe 450 ms, which is equal to reference duration (see Fig. 2.b, right), separately. The mean *RTP* of correct responses was highest in the gain group (*M* = .604, *SD* = .17), followed by the neutral (*M* = .594, *SD* = .15) and loss (*M* = .589, SD = .15) conditions. A one-way analysis of variance (*ANOVA*) using the Welch F-ratio showed that this difference was not statistically significant, F(.596, 15055) = 0, *p* = 1.0,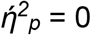. The median value for *RTP* of (450 ms) was highest in the loss group (*Mdn* = .593), followed by the neutral (*Mdn* = .580) and gain (*Mdn* = .577) groups. A Kruskal-Wallis test showed that this difference was not statistically significant, *H* = .01, *p* = .995, *ηH* = .04.

### Effect of reward on confidence

To examine the effect of the independent variable of reward on the dependent variable of confidence, we averaged the confidence scores of correct answers for each reward condition (gain, loss, and neutral) at six probe durations (300, 350, 400, 500, 550, and 600 ms), and the confidence scores of answers that subjects perceived to be longer on probe 450 ms, which is equal to the fixed reference duration (see Fig. 2.c). The median confidence score for correct answers was highest in the gain condition (*Mdn* = 4.51), followed by the loss (*Mdn* = 4.27) and neutral (*Mdn* = 4.09) conditions. A Wilcoxon signed-rank test showed that the difference between the gain and neutral conditions, *z* = -2.55, *p* = .0045 (one-tailed), *r* = -.42, and between loss and neutral conditions, *z* = -1.85, *p* =.033 (one-tailed), *r* = -.31 were statistically significant. The same test showed that the difference between gain and loss conditions was not statistically significant; *z* = -1.37, *p* =.182, *r* = -.23, (see Fig. 2.c).

Similarly, at the 450 ms, the median confidence for answers that were perceived longer was highest in the gain condition (*Mdn* = 3.79), followed by the loss (*Mdn* = 3.5) and neutral (Mdn = 3.29) conditions. A Wilcoxon signed-rank test showed that the difference between gain and neutral conditions was marginally significant; *z* = -1.63, *p* =.053 (one-tailed), *r* = -.27. The same test showed that the difference between gain and loss conditions, *z* = -.89, *p* =.134 (one-tailed), *r* = -.15, as well as between loss and neutral conditions z =..69, *p* =.25 (one-tailed), *r* =.2, were not statistically significant. This result indicates that the gain condition is perceived to last longer than the neutral condition with higher certainty. This pattern was even mirrored at the minimal perceptual level at a probe duration of 450 ms, although it reached a marginal level of significance (see Fig. 2.c).

### Effect of confidence on time perception in the reward context

#### Grouping individuals

In the previous sections, we found that time in the monetary gain condition was perceived as lasting longer than in the neutral condition, and that confidence was significantly higher in the monetary gain condition than in the neutral condition. However, from this data alone we cannot conclude that confidence influenced the perceived duration. To disentangle the role of confidence, we changed confidence to an independent variable by dividing subjects into two groups: high- (HC) and low- (LC) confidence. We classified subjects based on their overall mean confidence scores for the probe duration of maximum uncertainty (450 ms, probe equals reference duration) for the neutral condition (no gain and no loss). Please note that there are no correct answers for this probe, so we calculated the overall confidence level for both shorter and longer answers for the classification purpose.

We repeated all previous analyses on two groups of subjects and then compared them. To observe whether the subjects’ classification was reliable, we compared the confidence level of the perception of 450 ms (longer hits; subjects pressed longer choice) in the neutral condition. The HC group had higher confidence scores (*M* = 3.93, *SD* = 0.88) than the LC group (*M* = 2.82, *SD* = 0.6). An independent samples t test showed that this difference was statistically significant; *t*(14.07) = 3.12, *p* = .008 (see Fig.3.a). This pattern was repeated over other conditions as well, the HC group had higher confidence scores for gain (*M* = 4.52, *SD* = 0.76) than the LC group (*M* = 2.85, *SD* = 0.69). An independent samples t test showed that this difference was statistically significant, *t*(15.86) = 4.89, *p* < .001, and the HC group had higher confidence scores for the loss condition (*M* = 4.17, *SD* = 0.59) than the LC group (*M* = 2.9, *SD* = 1.02). An independent samples t test showed that this difference was statistically significant; *t*(12.78) = 3.24, *p* = .007 (see Fig. 3.a, left).

**Figure 3.**
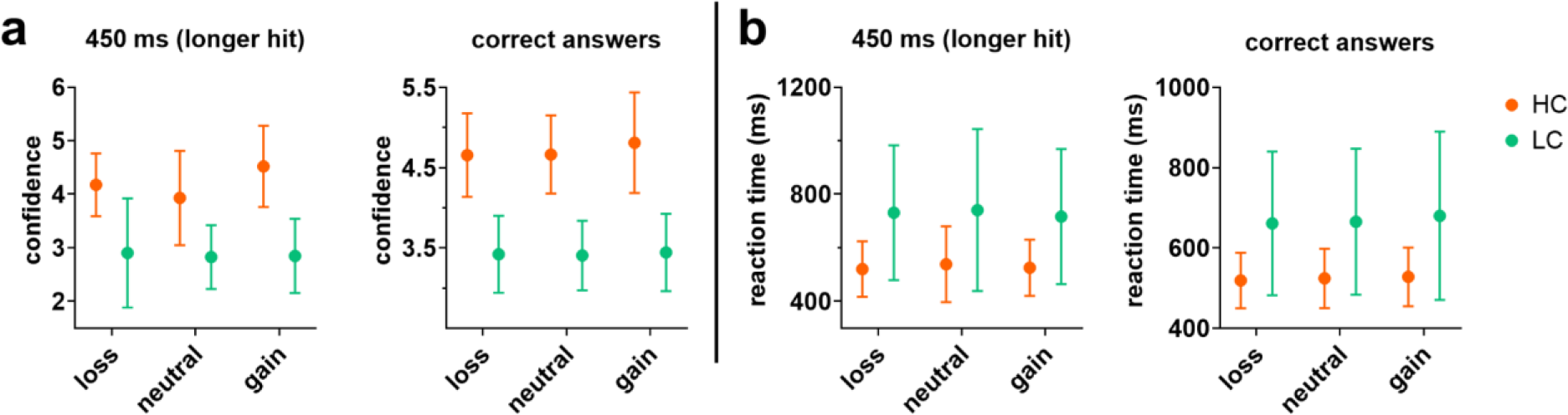
**a)** Group differences in confidence scores between LC and HC participants for 450 ms and correct answers. **b)** Group differences in reaction times between LC and HC participants for 450 ms and correct answers. (Note: all graphs are plotted based on the mean and standard deviation.)

The overall pattern was also replicated at the confidence scores for correct answers. The HC group had higher scores than the LC group for gain (*M* = 5.1, *SD* = 0.64 vs. *M* = 3.88, *SD* = 0.64); *t*(16) = 4.06, *p* < .001. for neutral (*M* = 4.93, *SD* = 0.57 vs. M = 3.67, SD = 0.43); *t*(14.92) = 5.26, *p* < .001, and for loss (*M* = 5, *SD* = 0.66 vs. M = 3.84, SD = 0.6); *t*(15.86) = 3.91, *p* = .001 (see Fig. 3.a, right).

We also compared *the RTP of* the two groups. *The* HC *group* responded faster than the LC group in all reward conditions and both correct answers and 450 ms (longer hits). The statistical test showed that all differences were significant *p* < .001. This indicates that the individuals are well split based on their minimal perceptual level in neutral condition (see Fig. 3.a and 3.b)^40^.

### Effect of reward on time perception within HC and LC groups

The previous section confirmed that the two groups were satisfactorily classified. To investigate whether reward influences time perception, we compared the *PSE*s between reward conditions in each group separately. In the HC group, *PSE* neutral had higher scores for outcome (*M* =.46, *SD* =.02) than *PSE* gain (*M* =.45, *SD* = .03). A paired-samples t test showed that this difference was statistically significant; *t*(8) = -3.57, *p* = .004 (one-tailed), *CI* [-.026, -.006], *Cohen’s dav* =.58 (see Fig. 4, left). Comparisons between (gain vs. loss) and (loss vs. neutral) revealed no differences. The *PSE* loss had higher scores for outcome (*M* = .46, *SD* = .03) than the *PSE* gain (*M* = .45, SD = .03). A paired-samples t test showed that this difference was not statistically significant; *t*(8) = - 1.58, *p* = .153, *Cl* [-0.025, 0.005], *Cohen’s dav* = .33. Similarly, *PSE* neutral had higher scores for the outcome (*M* = .46, *SD* = .02) than the loss (*M* = .46, *SD* = .03). A paired-samples t test showed this difference was not statistically significant; *t*(8) = -.76, *p* = .466, *Cl* [-.023, .012], *Cohen’s dav* = .21. In the LC group, *Wilcoxon signed-rank test* showed that comparisons between reward conditions were not statistically significant; *PSEs* (neutral vs. gain); *z* = -.18, *p* = .196 (one-tailed), *r* = -.04, (gain vs. loss); *z* = -.53, *p* = .346, *r* = -.13, and (neutral vs. loss); *z* = -1.01, *p* = .240, *r* = -.24. The overall pattern of the psychometric plot fitted to all subjects is replicated and similar to the pattern of the psychometric plot of high-confident individuals. This pattern is not replicated for low-confident subjects, meaning that the two groups perceived time differently and that only HC participants perceived the monetary gain condition as lasting longer than the neutral condition, not LC subjects.

**Figure 4.**
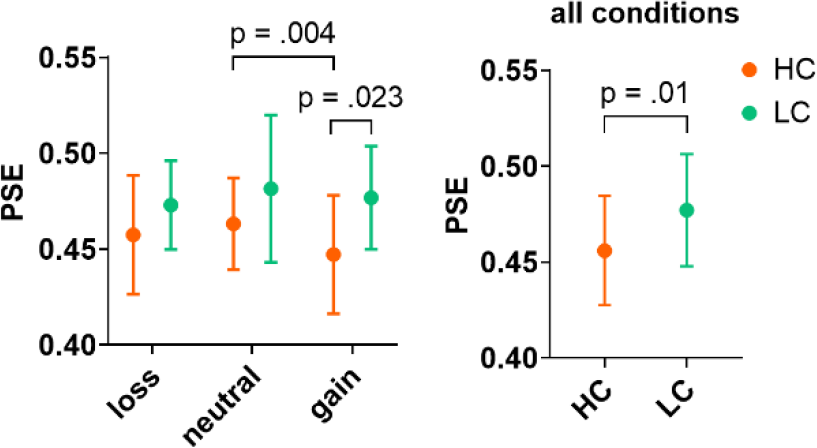
Comparison of *PSE*s between HC and LC groups across all reward conditions (left), Comparison of the mean of *PSE*s across all reward conditions between the HC and LC groups (right), (Note: all graphs are plotted based on the mean and standard deviation)

### Effect of reward on time perception between HC and LC groups

The LC group had higher scores for *PSE* gain (*M* = .48, *SD* = .03) than the HC group (*M* = .45, *SD* = .03). An independent sample t test showed this difference was statistically significant; *t*(15.71) = -2.17, *p* = .023 (one-tailed), *CI* [-.059, -.001] (see Fig. 4, left). The LC group had higher scores for *PSE* neutral (*M* = .48, *SD* = .04) than the HC group (*M* = .46, *SD* = .02). An independent sample t-test showed that this difference was not statistically significant; *t*(13.41) = -1.21, *p* = .123 (one-tailed), *CI* [-0.051, 0.014]. The LC group had higher scores for *PSE* loss (*M* = .47, *SD* = .02) than the HC group (*M* = .46, *SD* = .03). An independent sample *t-test* showed that this difference was not statistically significant; *t*(14.77) = -1.2, *p* = .124 (one-tailed), *CI* [-.043, .012]. These results indicate that subjects with high-confidence level perceived the duration of gain condition longer than subjects with low-confidence (see Fig. 4). In order to boost the statistical significance, we compared the mean *PSE* value of all reward conditions between two groups and we found the HC group perceived time longer than LC group (Fig. 4, right).

An independent sample *t test* showed this difference was statistically significant; *t*(15.71) = -2.17, *p* = .01 (one-tailed), 95% *CI* [-.059, -.001].

### Correlation and regression analyses

To examine the relationship between confidence and time perception in the monitory gain condition (condition of interest), *Pearson’s R* correlations (negative correlation) were performed between the *PSE* and confidence at the maximum uncertainty level (450 ms) and between the *PSE* and confidence of correct answers. The relationship between the *PSE* and confidence (450 ms) variables was found to be statistically significant; *r*(16) = - .57, 95% Cl[-.82, -.143], *p* = .0067 (see Fig. 5.a), as well as between the *PSE* and confidence at correct answers; r(16) = -0.44, CI[-0.752, 0.036], *p* = .034 (see Fig. 5.b). Nonsignificant results were found between the *PSE* and confidence variables in other conditions of. Additionally, a linear regression analysis was conducted to examine whether the confidence variable predicted *PSE* at the gain condition. The *PSE* gain condition was significantly predicted by confidence at the maximum uncertainty 450 ms (longer hits). The results of the regression model indicated that the confidence level of 450 ms explained 32.68% of the variance, and an overall collective significant effect was found; *R*^*2*^ = 0.33, *F*(1, 16) = 7.77, *p* = .013. The *PSE* gain condition was not significantly predicted by the confidence of correct answers.; *R*^*2*^ = 0.19, F(1,16) = 3.81, *p =* .069.

**Figure 5.**
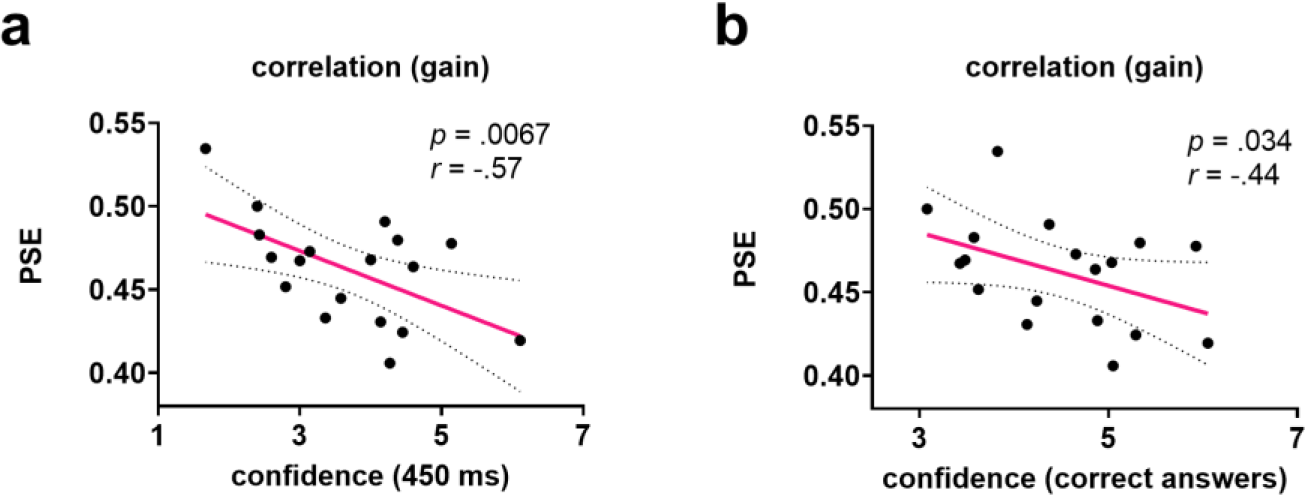
**a)** Pearson correlation (one-tail) between confidence (450 ms) and *PSE* in the gain condition **b)** Pearson correlation (one-tail) between confidence (correct answers) and *PSE* in the gain condition.

Moreover, further negative correlations were performed between RTP and confidence. The relationship between the RTP and confidence (450 ms, longer hits) variables was found to be statistically significant; r(16) = -.595, *p* = .015 (corrected for multiple comparisons). No significant results were found between other conditions (*p* > .05). This means that HC subjects were faster and more confident than LC in their responses.

### Early-late trials no effect on confidence

To determine whether there was a primacy or recency effect in the early trials on confidence scores, we split the data into two halves and compared confidence levels between the early and late halves. There were no significant results (*p* > .05) between the two halves of the data when all probe durations were considered (300 - 600, step = 50 ms), even with more stringent probe durations (400, 450, and 500 ms), (*p* > .05). This confirms that subjects performed the task well without fading the reference duration in their minds. In other words, if there was a significant difference between the confidence levels of early and late trials, a second experiment should have been designed and conducted in a different way, such as testing the reference and probes on a trial-by-trial basis, or by making the sessions shorter^41^.

## Discussion

In the current study we found a relationship between confidence, monetary reward, and time perception. The higher the degree of confidence, the longer time was perceived in a situation of financial gain compared to neutral and loss.

First, we replicated findings from previous studies showing that monetary reward has an effect on perceived time: the gain condition was perceived as lasting longer than the neutral condition. Evidently, the monetary gain condition had a smaller *PSE* than the neutral condition. This result is consistent with related studies that emphasized that modulations in time perception are due to the increased attentional deployment caused by the monetary gain condition^20,21^. They argued that high-money condition recruits subjective salience^42^ and therefore attracts more attention than low-or no-money. The greater increase in attentional deployment causes time to be perceived as extended. Arguably, in our study, the money loss condition had lower *PSE*s than the neutral condition, although they did not reach a significant level. This is inconsistent with studies that have found a positive relationship between arousal level and duration overestimation^43–45^ because reward processing studies have shown that anticipation of monetary loss, which facilitates avoidance behavior, involves the same arousal level as anticipation of monetary reward, which facilitates approach behavior^32^. The current results can be discussed as that monetary gain and loss^45^ conditions attract more attention from the valence gate than the arousal gate (robust reward processing studies, introduced that monetary incentives arising from the two orthogonal components, namely, arousal (from calm to excited) and valence (from pleasurable to aversive)), subject to the basic asymmetry in attentional choice between saving and earning money^46^. This suggests that altered time perception is based on specific mechanism of reward valence as one of the main components of Monterey Incentive Delay task, which may play an important role in the regulation of gaining behavior and decision making.

Next, we showed that the monetary reward influenced confidence. The monetary gain condition received a significantly higher confidence score compared to the neutral condition for correct answers and even for the probe 450 ms, albeit being marginally significant (*p* = .053). This finding is consistent with a study showing that participants were more confident in their decisions when learning to seek monetary gains than when learning to avoid monetary losses, despite equal difficulty and performance between these two contexts^27^. Neuroimaging studies^47,48^ have demonstrated that confidence is biased toward gain and seems to be beneficial for monetary payouts^49^. For example, a study demonstrated that confidence and difference in value are separate behavioral manifestations of the same underlying decision variable^50,51^. Also, a neurophysiology study found that decision certainty is encoded by the same neurons that reflect decision formation^28^, and in our study subjects were more confident while making decisions about the duration judgment of the gain condition compared to other conditions.

Third, we took an step forward and revealed how confidence levels affect time perception. We grouped subjects into high- and low-confidence individuals, and the findings remained unaffected for the high-confidence (HC) group. HC participants overestimated gain condition compared with the neutral condition. Previous studies showed that the correlation between confidence and objective performance varies for different people and is related to individual differences in brain structure^52^ and connectivity^53^, and to individual differences in mental calculation confidence^11^. Individual differences in confidence may also be related to differences in brain activity and neurochemistry. For example, studies have shown that high-confidence individuals show increased activity in brain regions associated with reward and motivation, such as the ventral striatum^47^. In contrast, individuals with low levels of confidence may exhibit reduced activity in these brain regions. Another possible explanation can be the regulatory focus theory that explains differences in strategic tendencies between individuals in their sensitivity to gains and losses, which result in variations in how they address problems^54^. Promotion-focused individuals have strong sensitivity to positive outcomes—gains and nongains. In contrast, prevention-focused individuals have a high sensitivity to negative outcomes—nonlosses and losses. To make an analogy, in our study HC subjects might be considered as individuals with a promotion focus and LC subjects as prevention focus, and this difference in their strategic tendencies causes a fundamental difference in their confidence level and subsequently the perceived time. This finding can be discussed by a very recent study that found a distinct pattern of inter-individual variations between individuals who used only perceptual differences to score their confidence and people who additionally used information that had no bearing on their discriminating judgments^30^. It is also possible to emphasize that our data imply that HC subjects made all decisions in goods space, and in value comparisons HC subjects valued more gain trials than other conditions^55,56^.

Finally, we found a negative correlation between confidence of correct answers and time perception and a regression between confidence of maximum uncertainty on 450 ms and time perception on gain condition across all subjects. This means that the more the subjects are confident in their correct answers, the longer the perceived time. Similarly, the more the subjects are confident in their errors, the longer the time perceived. Notably, the confidence in error trials that are perceived longer is predictive of time perception. This may be indicative of the metacognitive ability, and that in lower confidence is associated with higher metacognitive ability about knowing their errors ^57,58^.

In conclusion, individual differences in confidence levels can significantly influence how individuals perceive time in a monetary context. Understanding these differences is essential for developing effective strategies for motivating and engaging individuals in a range of settings, such as education, healthcare, and management. By identifying the factors that contribute to individual differences in confidence levels, we can develop tailored interventions that can enhance an individual’s motivation, focus, and ability to complete tasks successfully.

